# Engineering Genetically-Encoded Mineralization and Magnetism via Directed Evolution

**DOI:** 10.1101/085233

**Authors:** Xueliang Liu, Paola A. Lopez, Tobias W. Giessen, Michael Giles, Jeffrey C. Way, Pamela A. Silver

## Abstract

Genetically encoding the synthesis of functional nanomaterials such as magnetic nanoparticles enables sensitive and non-invasive biological sensing and control. Via directed evolution of the natural iron-sequestering ferritin protein, we discovered key mutations that lead to significantly enhanced cellular magnetism, resulting in increased physical attraction of ferritin-expressing cells to magnets and increased contrast for cellular magnetic resonance imaging (MRI). The magnetic mutants further demonstrate increased iron biomineralization measured by a novel fluorescent genetic sensor for intracellular free iron. In addition, we engineered *Escherichia coli* cells with multiple genomic knockouts to increase cellular accumulation of various metals. Lastly to explore further protein candidates for biomagnetism, we characterized members of the DUF892 family using the iron sensor and magnetic columns, confirming their intracellular iron sequestration that results in increased cellular magnetization.

## Introduction

Inorganic nanomaterials have been used in a wide range of biological applications including fluorescent or plasmonic labelling for imaging, magnetic labelling for extraction and high throughput sequencing and drug-delivery^1–3^. However, unlike genetically-encoded labels such as green fluorescent protein (GFP), chemically synthesized inorganic nanomaterials, despite their versatile physical and chemical properties, are ultimately limited in their biological application by their lack of integration with the genetic circuitry of the cell. Synthetic biology can bridge this gap by programming cells to controllably synthesize their own nanomaterials in response to biological signals. Those nanomaterials can be further tailored within cells to interact with other components and transduce biological signals downstream.

There are few examples of bio-synthesized inorganic nanomaterials in Nature. Certain species of bacteria and archaea can mineralize nanoparticles via proteins or metabolites that reduce toxic metal cations^1,4^. Notably, magnetotactic bacteria of the genus *Magnetospirillum* naturally synthesize crystalline magnetite nanoparticles and align them as a passive navigation compass for the cell in its natural environment^5–7^. Despite speculation on the presence of similar inorganic magnetic nanoparticles in animals such as fish and humans, no such biomineralization pathways have been confirmed so far^8–11^. However, all cells do use inorganic bio-mineralization to maintain near constant concentrations of essential trace metals via high affinity chelators and storage proteins for times of excess. One prominent example are the ferritins, a ubiquitous class of proteins found in all domains of life that play a crucial role in iron homeostasis^2,12–18^. Ferritins form shells composed of 24 monomers each, creating an inner cavity in order to store iron in a hydrated amorphous form of iron oxide similar to the mineral ferrihydrite. (Figure 1a, b) Iron oxide is biocompatible and magnetic depending on its crystal structure. However, the mineralized iron stored inside natural ferritins exhibits poor crystallinity which facilitates iron release in times of need but also results in a very modest inherent magnetic moment^19–21^. Even though the factors that control crystallization and hence the properties of the magnetic nanoparticles inside ferritin cages are not completely clear, natural ferritins still represent an excellent starting point for protein engineering aimed at increasing the inherent magnetism of ferritin particles. Engineering increased bio-magnetism would open up the way for advanced applications in non-invasive biological sensing, imaging and actuation^22–35^. Here, we employ directed evolution of ferritin to enhance the magnetism and biomineralization capability of engineered *E. coli*. In addition, a novel genetic biosensor for cytoplasmic free iron was developed to easily measure biomineralization efficiency *in vivo*, combined with genetic manipulations of metal transporters to optimize intracellular metal levels^36–40^. (Figure 1c)

**Figure 1.**
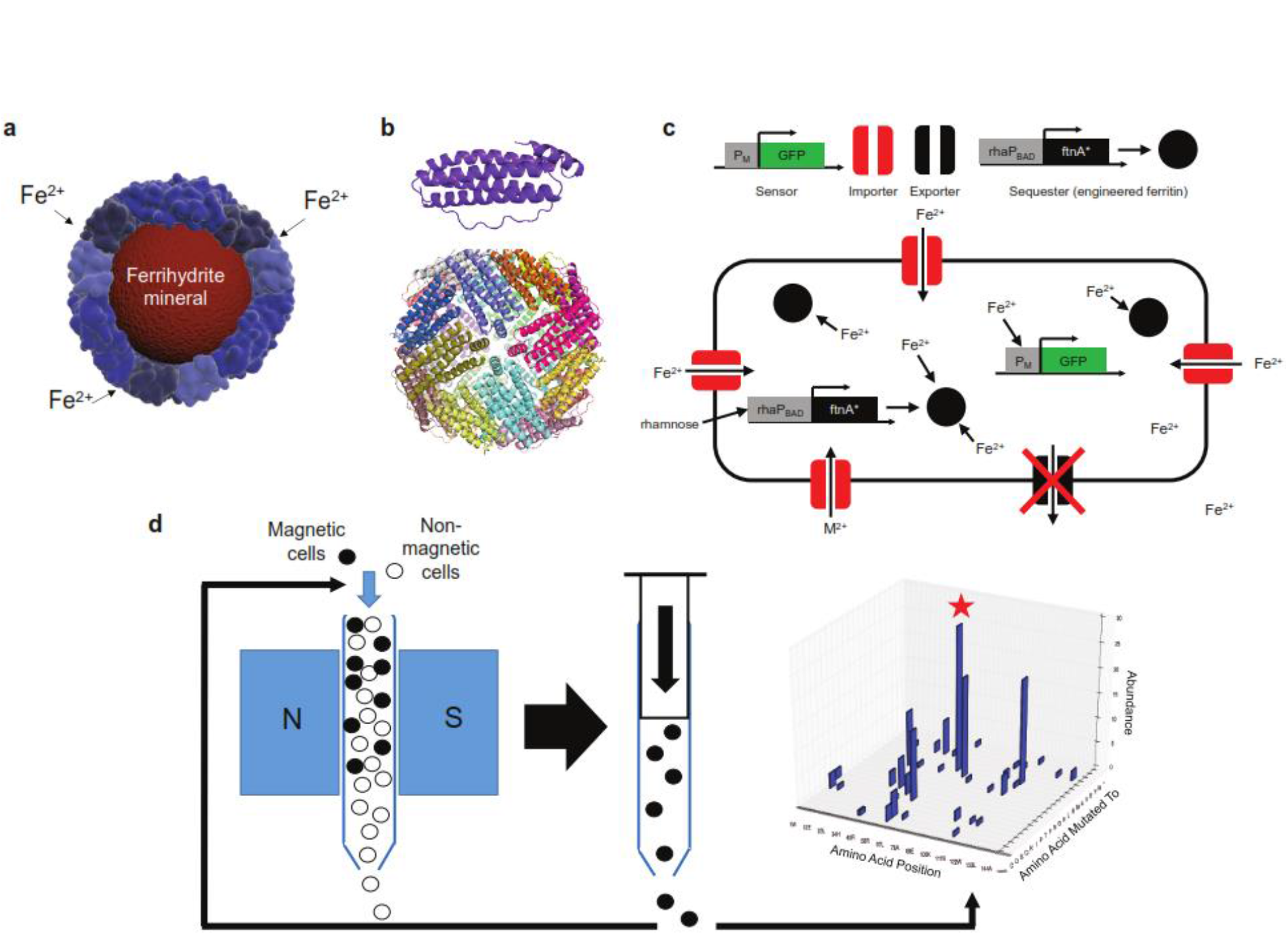
Engineering cellular magnetism. **(a)** Schematic of the ferritin protein shell encaging a ferrihydrite nanoparticle **(b)** crystal structure of the ferritin monomer (top) and the self-assembled, 24-homomers cage (bottom). **(c)** The cell engineered to accumulate metals by knockout of genomic exporters (black) and expression of importers (red). Mutant ferritins particles (black spheres) induced by a rhamnose promoter biomineralize iron into intracellular magnetic particles. A genetic fluorescence sensor monitors intracellular free Fe^2+^ level. **(d)** directed evolution for increased magnetism: iterative selection by high gradient magnetic column of a library of cells expressing randomly mutated ferritins was carried out over 10 days (1 cycle/day). Subsequent sequencing analysis of the magnetically retained mutants enabled discovery of mutations in key residues (e.g. red star: T64) that enhanced cellular magnetism.

## Results

We discovered novel mutations in ferritin that increase cellular magnetization via "magnetic evolution", combining random mutagenesis with iterative magnetic column selection (Figure 1d). Specifically, a randomized library of *E. coli* ferritin (*ftnA*) mutants was cloned and transformed into *E. coli*. Magnetic cells from this population were enriched through ten iterative rounds of growth followed by passage through high-gradient magnetic columns. Eventually, the selected population of cells was plated and sequenced, and the most frequent mutations were identified and validated in freshly cloned and transformed cells (Figure 2a). We further combined and tested additional mutants informed by results of the initial random mutagenesis experiment. In total, we tested 38 mutants of *ftnA* via overexpression in *E. coli*. (Table S1) The native ferritin-clan genes (*ftnA, bfr, dps*) were knocked-out to prevent interference with the magnetic evolution approach. Changes in cellular magnetism were compared via measurement of the retention fraction of cells in high-gradient magnetic columns. The results show that the majority of mutants, compared to wild-type ferritin, significantly increase levels of magnetism of cells (Figure 2b). The top mutant (m1) contains the double point mutations H34L and T64I at the 2-fold symmetrical "B-type channel" (Figure 2c). These point mutations result in smaller or less polar residues occupying the entrance to the "B-type channel". This would likely increase the size of the channel encouraging ion diffusion into the core^41^. The two sites were individually mutated to alanine and demonstrated increased magnetism compared to wild-type while still being less magnetic than the best double mutant. Furthermore, a BSA-calibrated SDS-PAGE gel of the cells shows that the mutants, in particular the most magnetic ones, exhibited reduced protein expression compared to wild-type, confirming that the increased cellular magnetism is not caused by increased levels of ferritin expression. (Figure 2d, S1)

**Figure 2.**
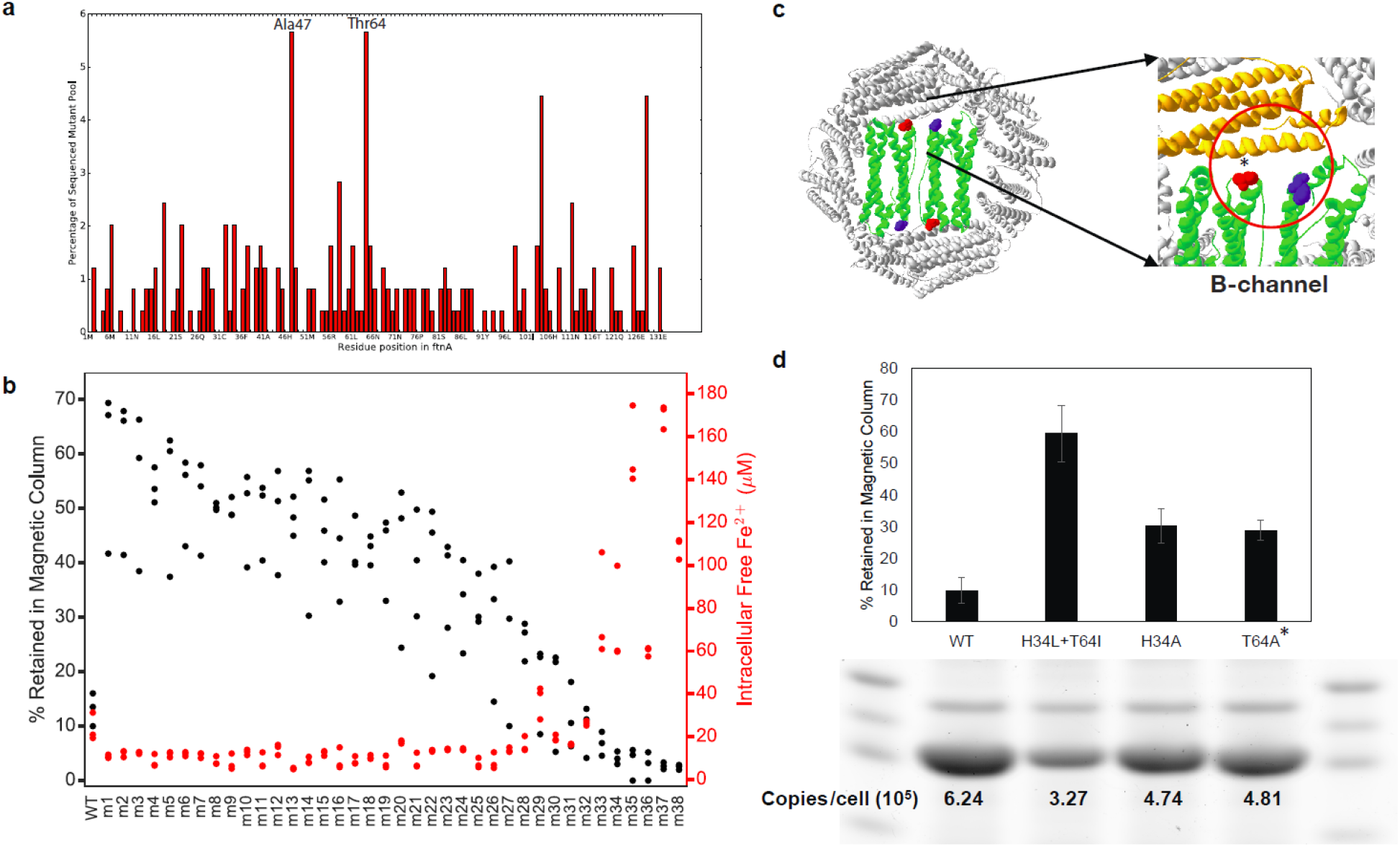
Mutations in ferritin proteins enhance magnetism. **(a)** Histogram of mutations from sequencing samples of the magnetically-evolved culture reveals mutations of key residues (e.g. Thr64, Ala47) that enhance biomagnetism. **(b)** Magnetic column retention levels (black dots) for all engineered ferritins (Table S1) including H34L+T64I (m1), sorted in descending order of their observed averages. Most mutants show significant increase over wildtype (m1−m21, p<0.05 by two-tailed t-test, N=3). The intracellular free iron concentrations inferred from genetic iron sensor measurements (red dots) are anti-correlated to magnetic column retention levels. **(c)** The most magnetic ferritin mutant (H34L+T64I) is doubly mutated at the "B-channel" (red circle) that transports iron. **(d)** The importance of the His34 (blue) and Thr64 (red) sites are individually validated by single mutants, demonstrating significant magnetic retention levels over the wildtype (WT). Furthermore, quantitative SDS-PAGE gel analysis (cropped to show ftnA band, full-length gel is presented in Supplementary Figure S1) demonstrates weaker expression for the mutants compared to the wildtype, confirming that increased magnetism is not caused by mere increases in protein and nanoparticle quantities.

We further developed a genetic fluorescent iron sensor that demonstrates an increase in cytosolic iron sequestration in strains expressing mutant ferritins with increased magnetism. The iron sensor employs the *E. coli* promoter *fiu*^42^, which contains four overlapping binding sites for the transcription factor *fur* (ferric uptake regulator). *Fur* bound to Fe^2+^ represses downstream expression of GFP on a low-copy plasmid (p15A origin) (Figure 3a). Hence the *fiu*-based sensor reports depletion of free iron in the cytoplasm. Further calibration using a range of concentrations of the cell-permeable ferrous iron chelator bipyridine (bpd) allows conversion of culture density-normalized fluorescence to free iron concentrations per cell (Figure S2). This converted concentration was monitored over time for *E. coli* growing from exponential to stationary phase in LB medium supplemented with different Fe(II) sulfate concentrations (0 to 5 mM) over a range of ferritin induction levels (0% to 0.2% rhamnose). The time courses show rapid sequestration of intracellular iron under high ferritin inductions and decreased time to reach peak iron concentrations (Figure 3, b-g). Without induction, significantly higher intracellular Fe levels were observed when iron was supplemented into the medium at 5 mM, close to the observed viability limit for *E. coli* (∼10mM). Comparing the final readout at 15 h between cells expressing wild-type and the most magnetic mutant (m1) shows that the mutant sequesters more iron, especially at low induction levels below 0.01% rhamnose (Figure 3, h-I, replicated in Figure S3, S4). We also found that for the ferritin mutants, intracellular iron concentration in stationary phase is generally anti-correlated with magnetic column retention (Figure 2c). Considering the lower protein expression levels of the mutant as estimated quantitatively from SDS-PAGE gel analysis of whole-cell lysates (Figure 2d), iron sequestration (i.e. decrease of iron concentration) generally correlates with greater magnetic retention.

**Figure 3.**
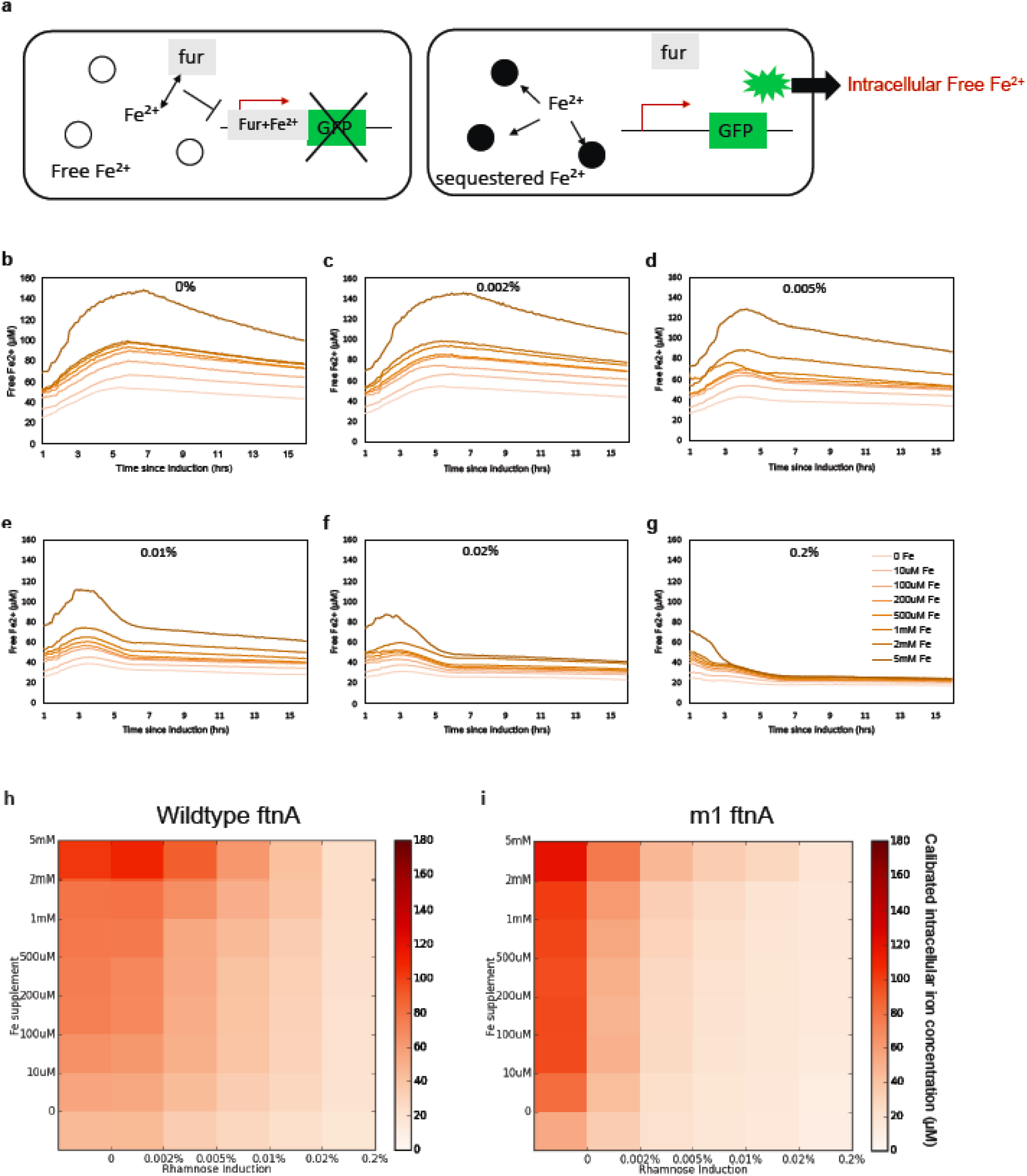
Genetic fluorescent sensor monitors cellular Fe^2+^ sequestration in WT and mutant ferritin expressing cells. **(a)** Free Fe^2+^ binds to the ferric uptake regulator (apo-fur). The Fe-bound fur binds the fiu promoter sequence to repress transcription of GFP (right). Sequestration of free Fe^2+^ by ferritins increases sensor fluorescence which with calibration is converted to the intracellular free Fe^2+^ concentration. **(b-g)** Calibrated free Fe^2+^ concentrations in *E. coli* expressing wildtype ferritin from up to 15 hours after induction by 0% (b), 0.002% (c), 0.005% (d), 0.01% (e), 0.02% (f), and 0.2% (g) rhamnose with media Fe^2+^ supplement of 0 μM, 10 μM, 100 μM, 200 μM, 500 μM, 1 mM, 2 mM, 5 mM. Without ferritins high media supplement at 5mM can dramatically alter the intracellular iron homeostatic setpoint. At high induction levels free Fe^2+^ is efficiently sequestered up to the highest media supplement concentration. The heatmaps with color saturation proportional to calibrated intracellular free iron levels show that compared to the wildtype **(h)**, the best ferritin mutant **(i)** is much more effective at sequestering iron at lower protein levels (0.01% rhamnose induction) and at high environmental iron concentration (up to 2 mM). This is consistent with their greater magnetism despite lower protein expression.

Cellular magnetism and mineralization were further confirmed by a range of characterization methods. The increased magnetic moment of the mutant cells was visualized directly by growing cells in LB rich medium in titer plate over ring-shaped permanent magnets^21^. Over-expression of mutant ferritins produced a sharper "image" of the magnets compared to wild-type expression, due to greater cellular magnetic moment and hence force on the cells from the surface field gradient of the permanent magnets (Figure 4a). Iron-loaded ferritins could be visualized in some cell cross-sections as aggregates of electron-dense puncta using transmission electron microscopy (TEM) (Figure 4b). Due to their paramagnetism, the local field inhomogeneity around the particles in the presence of an applied magnetic field increases spin-spin relaxation of the spins of protons in surrounding water molecules, producing contrast for T2^*^ based MRI using a gradient-echo pulse sequence. The significant reduction in T2^*^ was only observed for over-expression of the mutant m1, attributable to the much greater moment of the individual particles (Figure 4c). Lastly, Superconducting Quantum Interference Device (SQUID) measurement of cellular magnetic moment shows that only overexpression of the mutant ferritins leads to a positive paramagnetic contribution over the negative diamagnetic contribution from cells (Figure 4d).

**Figure 4.**
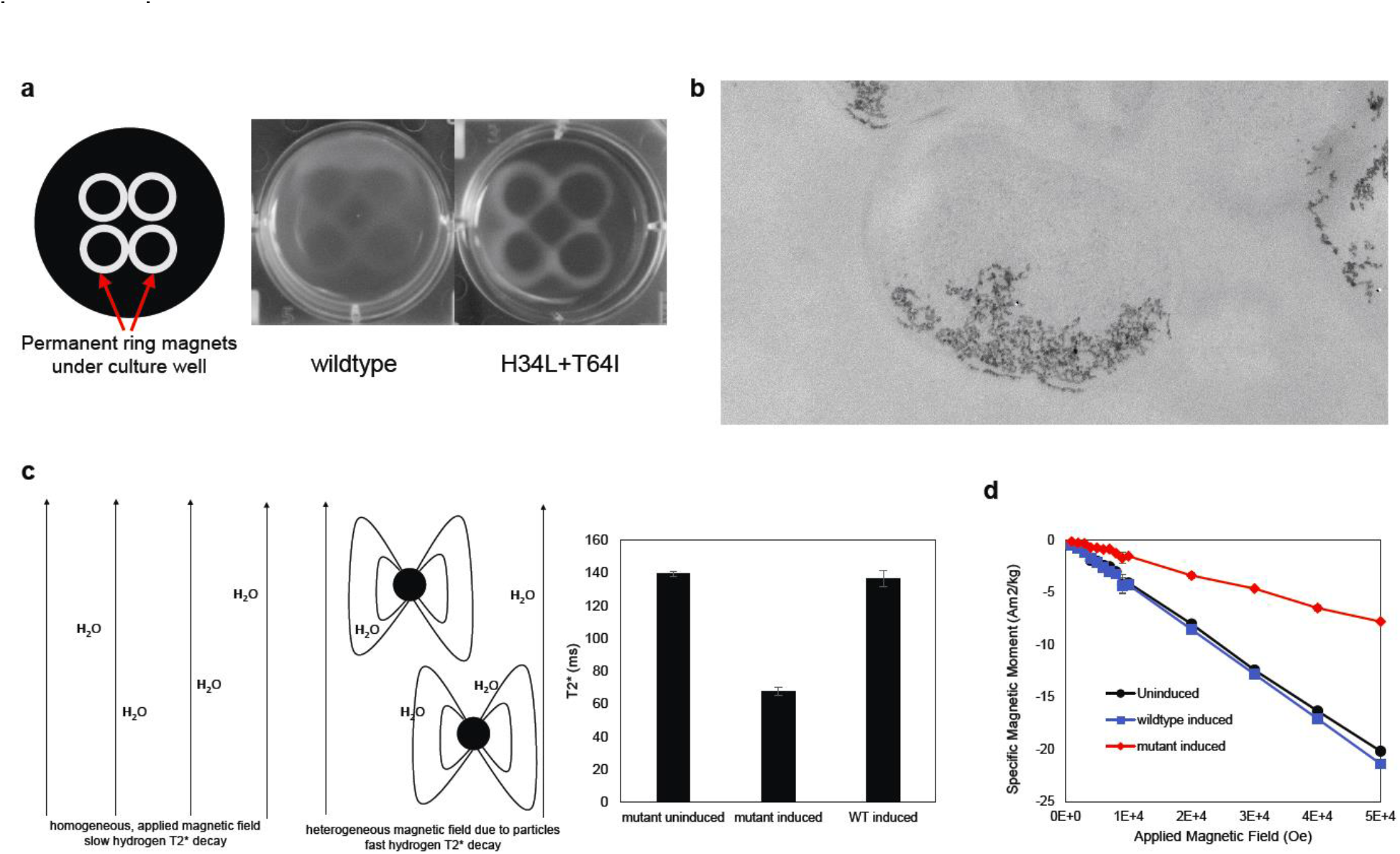
Characterizing cellular magnetism and biomineralization. **(a)** In liquid culture, *E. coli* expressing the mutant are attracted to ring-shaped magnets much more than the wildtype, due to stronger magnetic moments. **(b)** TEM images show electron-dense magnetic particles in a cross section of fixed and embedded cells that expressed engineered ferritins. **(c)** In MRI imaging, the presence of magnetic ferritin particles creates local magnetic field inhomogeneities that variably alter nearby proton relaxation rates and accelerate T2^*^ decay. Triplet cultures of E. coli expressing ferritin mutants more significantly decreased T2^*^ relaxation time compared to both no expression or overexpressing the wildtype (p<0.05 by two-tailed t-test, N=3), which creates imaging contrast. **(d)** SQUID magnetometry shows that only expressing the mutant ferritin led to positive paramagnetic increases in the cellular magnetic moment relative to the uninduced control.

In addition to iron-sequestration, the ferritin mutants have potential to co-sequester other elements. We first engineered *E. coli* with knockouts in three divalent cation exporters (*rcnA, fieF, zntA*) and also the iron master regulator *fur*, causing increased metal accumulation in the cells, particularly for iron, zinc and cobalt (Figure 5). Knockout of *zntA* alone seemed sufficient for increasing zinc concentrations, particularly when ferritins were over-expressed. However, due to compensatory effects among the exporters and also *fur* regulation, only knocking out *fur* along with the three exporters (*rcnA, fieF, zntA*) significantly increased cellular iron levels as measured by inductively coupled plasma mass spectrometry (ICP-MS). We also found co-expressing importers could increase iron levels and cellular magnetism (data not shown). We then tested several elements known to absorb to or alloy with iron oxide: cobalt, nickel, cadmium and arsenic. Cells were exposed to varying concentrations of these elements. Only strains expressing magnetic mutant ferritins exhibited a decreased growth defect, suggestive of toxic metal sequestration by over-expressed ferritin mutants (Figure S5). The benefit of expressing the mutants over the wild-type ferritins may be explained by the increased biomineralization rate of the mutant ferritins which aids in co-sequestration of other elements into the growing iron oxide core.

**Figure 5.**
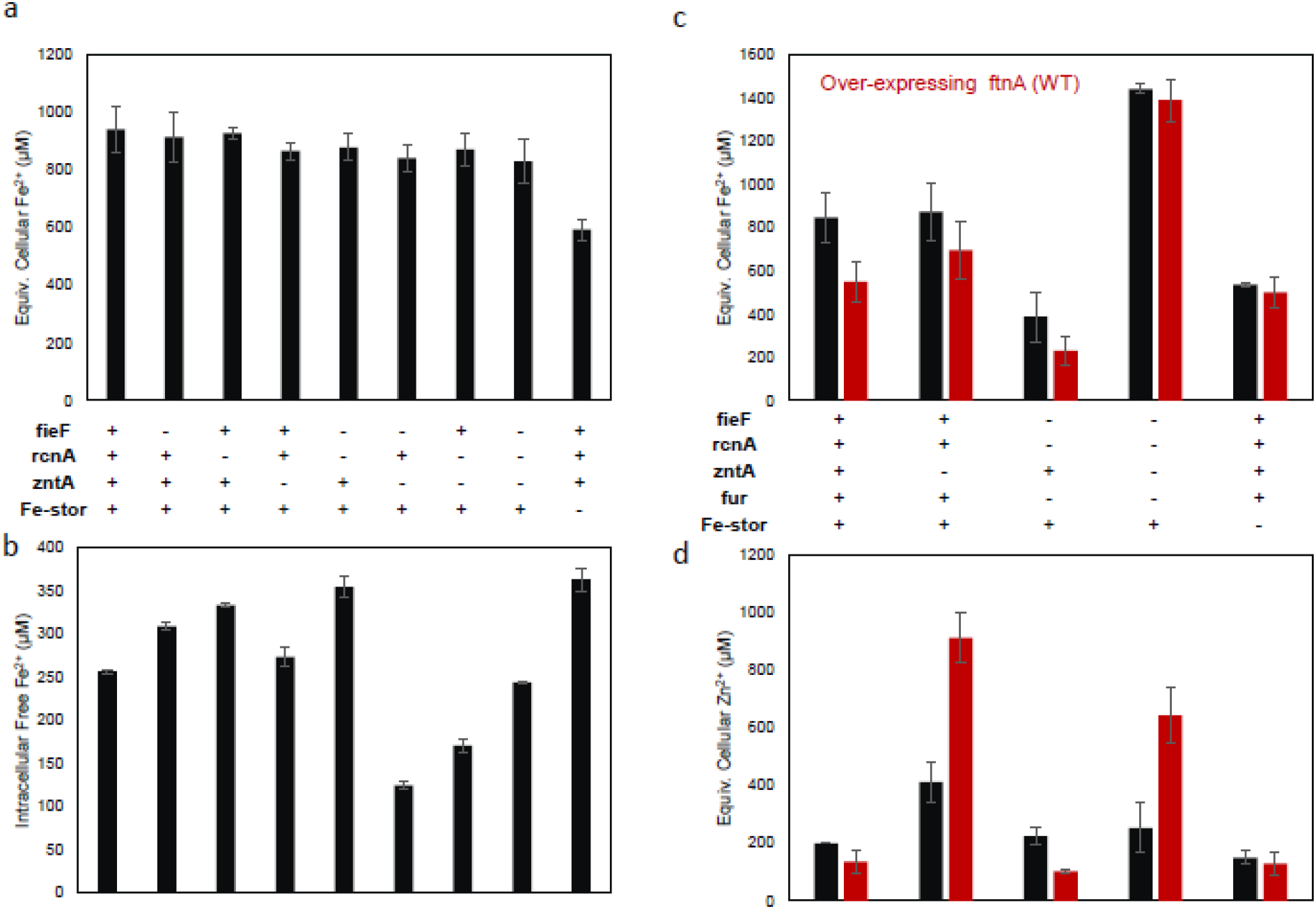
Effect of genomic knockouts on cellular metal concentrations. **(a)** ICP-MS measurement of Fe concentration for combinations of ferritin or metal exporter knockouts (−). With the master regulator fur protein present, iron homeostasis is maintained unless multiple ferritin homologs are knocked out (Fe-stor = *ftnA*, *bfr* and *dps*). **(b)** Genetic Fe-sensor measurement of intracellular free Fe^2+^ concentration shows effect of knockouts in the same strains despite homeostasis of total Fe in A. **(c)** ICP-MS of total Fe concentration for additional mutants with *fur* KO, which increases total Fe level. Over-expressing ftnA (red bars), however, did not increase iron level. **(d)** ICP-MS of zinc concentration for additional mutants with *fur* KO. Knockout of *zntA* alone dramatically increases cellular Zn levels, especially when expressing ferritins capable of binding zinc. *zntA* plays a crucial role in Fe export in the absence of the iron exporters and regulator *fur*, resulting in dramatically decreased Fe levels in contrast to its knockout. *zntA* knockout in combination with either *fieF* or *rcnA* KO further increases intracellular Fe sequestration. Comparison of total Fe content (a) vs. free Fe^2+^ via the genetic sensor (b) illustrates the interplay between metal transport and storage in cells.

Lastly to explore constructs other than the well-known ferritins as candidate starting points to modulate biomagnetism, we studied the iron sequestration and magnetic properties of Domain of Unknown Function 892 (DUF892) proteins, which fold into a similar 4-helix bundle as ferritins and are thus distantly related in structure-based phylogeny^43^ (Figure 6a). Three predicted, uncharacterized proteins in the DUF892 family (A0A072C8A3, F7XJ33 and Q1QGY5) were cloned and over-expressed in wildtype *E. coli* containing the fluorescent iron sensor. The converted intracellular iron levels based on fluorescence measurements were markedly decreased compared to the no-expression control, with a concomitant increase in the percentage of cells retained in magnetic columns suggesting biomineralization (Figure 6b). These results confirm that the function of DUF892 proteins is likely similar to that of ferritins, namely to sequester free iron in times of excess or stress.

**Figure 6.**
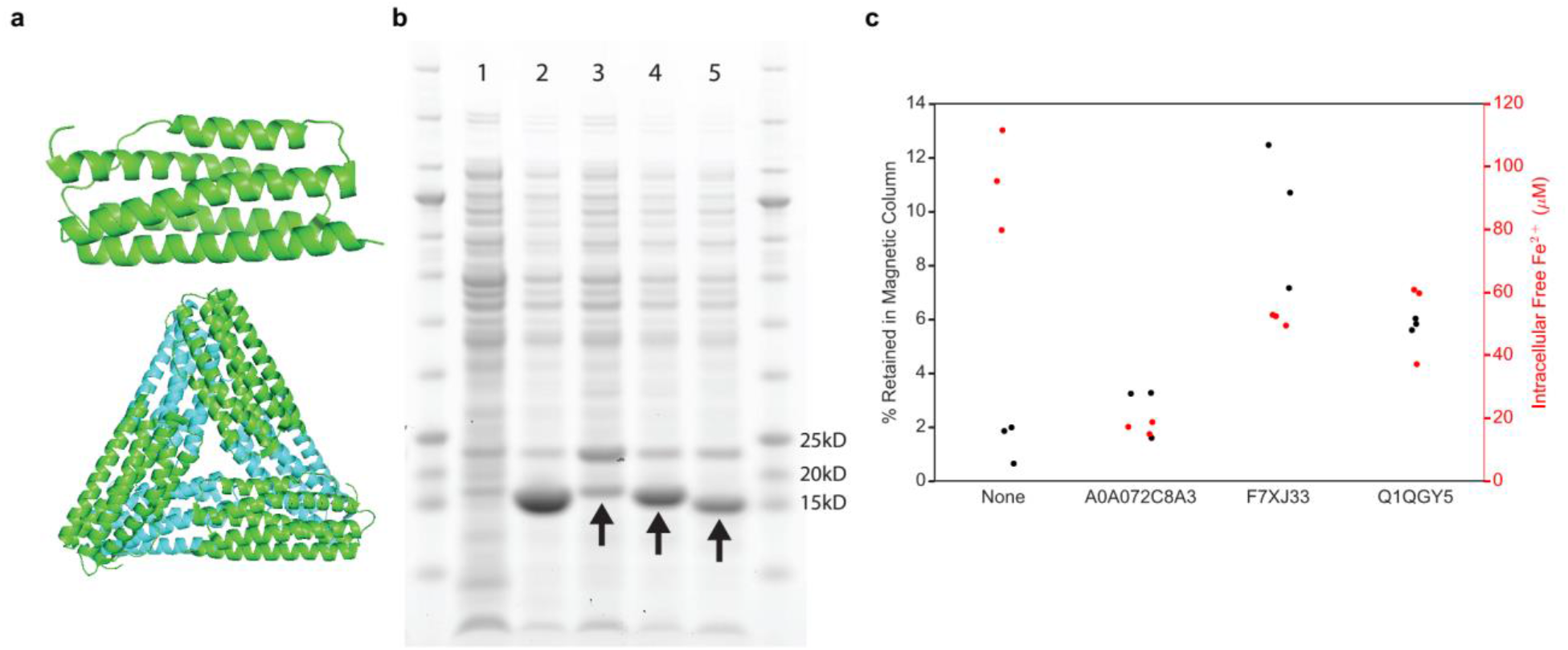
DUF893 proteins sequester iron and increase cellular biomagnetism. **(a)** The crystal structure of a DUF893 family protein monomer (*E. coli* YciF; PDB ID: 2GS4) shows a similar 4-helix bundle as ferritin (top), but with different predicted quaternary structure (bottom). **(b)** Verification by SDS-PAGE of cell lysates for the over-expression in *E. coli* of three predicted proteins in the DUF893 family: A0A072C8A3 (lane 3), F7XJ33 (lane 4), Q1QGY5 (lane 5). Lane 1 is no-expression control. Lane 2 is over-expression of ftnA. The over-expressed proteins have similar theoretical molecular weights with bands running close to 15kD mark (arrows). The common band close to 25kD mark represents GFP output of the iron sensor integrated into the E. coli **(c)** The new DUF893 proteins demonstrate significantly decreased intracellular free iron levels (p<0.05 by two-tailed t-test, N=3) and increased level of cellular magnetism by magnetic column retention measurement compared to the no-expression control.

## Discussion

In this study, we used directed evolution by magnetic selection to discover point mutations in ferritin that enhance iron sequestration and the magnetic phenotype of cells expressing ferritin mutants. Increased cellular magnetism was shown to lead to physical attraction to magnetic field gradients and improved MRI contrast. We further developed a new genetic iron sensor in *E. coli* that fluorescently detects variations in intracellular free iron levels to validate iron sequestration by ferritin mutants and other previously uncharacterized iron-sequestering proteins.

The genetic iron sensor was used to demonstrate the ability of magnetic ferritin mutants to more efficiently sequester iron. In addition, it revealed several features of cellular iron homeostasis. First, intracellular free iron concentration of *E. coli* increases sub-linearly with increasing iron supplement in culture media until around 2 mM, above which the sensor displays a sudden increase in intracellular free iron levels, coincident with significant growth defect and death (Figure 3b). However, over-expression of ferritins rapidly sequesters extra iron, independent of the range of supplement concentrations tested, to yield a low, steady iron level in the cells. The lower free iron level under ferritin expression despite high supplement levels for iron indicates that the cells are not able to efficiently compensate for lost iron by increasing import. This is due to the active nature of iron import relying largely on the synthesis of iron siderophores which may be the bottleneck in maintaining iron homeostasis during rapid iron starvation.

The directed evolution and magnetic selection of ferritins is an efficient strategy to obtain magnetic protein constructs. The mutations found in the most magnetic ferritin at the ferritin B-type channel directly affect iron transit into the protein shell. It is commonly recognized that the 3-fold symmetrical channels of the ferritin particle are most important for iron transit. It has also been assumed that iron entry requires Fe(II) to first localize to and oxidize at the ferroxidase active site close to the center of the 4-helix bundle. The role of the B-type channel has not been highlighted until recently^41^. Here, we show mutations around the B-type channel directly affect iron sequestration rates *in vivo*. In particular, the H34L and T64I mutations likely enlarged the B-type channel to increase the influx of un-oxidized Fe(II) ions. Histidine has a larger side chain compared to leucine. On the other hand, threonine at position 64 could serve as a C-terminal cap of the second major helix via hydrogen bonding, hence its replacement could change pore structure to favor ion entry. This in turn may affect not only the concentration but also the balance of Fe(II) and oxidized Fe(III) species in the core. Natural ferritin mineralizes mostly Fe(III) ions into Fe_2_O_3_, which may exhibit antiferromagnetic coupling of iron atom spins resulting in a very low magnetic moment. However, magnetite (Fe_3_O_4_), the most magnetic form of iron oxide, contains mixed valence atoms of Fe(II) and Fe(III). Hence the influx of Fe(II) into the ferritin core without oxidation could change the stoichiometry of iron valencies inside the nanoreactor to favor mineralization of the more magnetic magnetite. The increased iron-sequestration and cellular magnetism of mutants resulting from the directed mutation of residues His34 and Thr64 at the B-channel to alanine supports this hypothesis. Interestingly, we found that certain mutants with N-terminal "SpyTag" also increased magnetization (Fig. 2, Table S1). SpyTag was introduced to allow potential covalent attachment of ferritin particles onto other targets possessing the corresponding "SpyCatcher" tag^44^. Unlike the H34L and T64I double mutant, however, Spy-tagged ferritins were present in higher amounts suggesting an increase in the number of ferritin particles per cell. Furthermore, the extra peptide at the ferritin N-terminus, close to the 2-fold symmetric B-type channel in the self-assembled particle, could alter local structures around the pore to affect its properties and potentially increase iron flux.

Improving biomineralization and biomagnetism can lead to diverse applications. Cells genetically modified to accumulate and store toxic or valuable metals can enable cell-based bioremediation or mining. On the other hand, the magnetic particles from ferritin mutants could serve as noninvasive, genetically-encoded reporters of biological activity in deep tissues transparent to magnetic fields, as demonstrated by their enhanced MRI relaxation rate in T2^*^ MRI. T2 imaging using spin-echo sequence is more commonly reported for magnetic particles. However, spin-echo sequences are designed to cancel the dispersion in transverse relaxation due to magnetic field inhomogeneity produced by the ferritin iron-oxide particles in the static limit. Hence the T2 signal change is dependent not only on the magnetic moment but also on change in the particles' local magnetic environment due to diffusion during measurement. Hence T2 measurement is likely dependent on measurement parameters like sequence duration, whereas T2^*^ gradient echo is not. For *in vivo* imaging, both T2 and T2^*^ imaging modes can be used to detect contrast generated from endogenous magnetic particles. The ferritin mutants could allow noninvasive reporting of *in vivo* biological signals from engineered cells such as bacteria in the gut, or immune cells targeted to cancer tumors. There are further potential application of engineered intracellular magnetic particles as magnetic force or magneto-thermal actuators to manipulate proteins such as ion channels or subcellular compartments to control cellular behavior. Such applications include control of calcium-based signaling in mammalian cells, particularly neurons to induce or inhibit action potentials (e.g. "magneto-genetics")^33,34^, and manipulations of subcellular organelles and structures (e.g. magnetic tweezer). However, the natural size of the ferritin cage, at 8 nm in core diameter, is too small to create sufficient magnetic moment from the encaged nanoparticle to achieve desired levels of sensitivity and reliability in most actuation applications^19^. Larger protein shell templates are desired, and Nature may contain several such templates such as encapsulins^45^. The approach developed here of using a fluorescent metal sensor and magnetic directed evolution could be similarly used, such as done to characterize the function of DUF892 proteins here, to discover additional novel candidates that would enable enhanced biomagnetism applications in mammalian biology.

## Methods

### Construction of *E. coli* knock-out strains

We chose *E. coli* BW25113 as the background to introduce plasmids expressing recombinant proteins or conducting sequential knock-out of genes via P1 transduction from knock-out strains in the KEIO collection of non-essential *E. coli* knock-out strains. Plasmid pCP20 was electroporated into the transduced cells and grown at 37°C on LB agar plate to induce expression of recombinase to that flip out the Kanamycin resistance cassette used for transduction selection, as well as loss of the pCP20 plasmid. The genes knocked out include genes that serve as endogenous iron storage proteins (ftnA, bfr, dps and their combinations), genes that serve as exporters of metal cations (fieF, rcnA, zntA and their combinations) and the iron master regulator fur. BW25113 with all ftnA, bfr and dps served as the background strain for expressing recombinant ferritins so as to minimize the small background of endogenous iron storage proteins. BW25113 with all fief, rcnA, zntA and fur knockous served as the metal over-accumulation strain to minimize metal export and down-regulation of import by fur in a metal-rich environment.

### Construction of recombinant ferritin

*E. coli* ferritin gene ftnA was cloned into a high copy-number plasmid (pUC Origin of replication) with rhamnose inducible promoter^46^ (*rhaP_BAD_*, with native *E. coli* transcription factors RhaS and RhaR) and kanamycin resistance cassette via Gibson Assembly^47^. The DNA plasmid was verified by Sanger Sequencing (Genewiz) and transformed into *E. coli* BW25113 cells via electroporation. Expression of ftnA was induced in cells by adding rhamnose to cell culture (maximum 0.2%) during log-phase growth (OD600∼0.4).

### Directed evolution of magnetic ferritin mutants

Error-prone PCR (Agilent GeneMorph II Random Mutagenesis Kit) was used to generate random mutants of the *E. coli* ferritin (ftnA) sequence with on average one mutation per copy. The PCR products were cloned into the vector with rhamnose promoter and transformed via electroporation into *E. coli* strain containing the GFP-based genetic iron sensor and triple knockout of endogenous iron sequesterers (ftnA, bfr, dps). This library of cells each expressing a particular ftnA mutant was grown and induced in log-phase with 0.2% rhamnose to initiate high level expression of ferritins and simultaneously exposed to 100uM iron (II) sulfate as supplement. The cells were cultured with good oxygenation in deep-well plates shaken at 900rpm. In saturation phase (∼20 hours post metal-exposure), the cells were filtered through the magnetic column (Miltenyi LD column) placed between neodymium permanent magnets (K&J Magnetics Inc., BX8C4-N52) to create a high magnetic field gradient that help to retain cells that are more magnetic in the column whereas the cells expressing less magnetic ferritin mutants are flushed out. Subsequently, the permanent magnets are removed and the previously retained magnetic cells were eluted separately, grown again to log-phase, and re-induced and exposure to iron in fresh media to iterate another day of growth and magnetic selection. In total, 10 days or iterations of magnetic selection were conducted before the final eluted magnetic cells were plated. Around 100 colonies were picked and sequenced by Sanger sequencing (Genewiz). A Python script was used to analyze the sequences to determine the most frequent or representative mutations. The top mutations were re-constituted in wildtype ftnA plasmid (NEB Q5 Site-Directed Mutagenesis Kit) and verified for increased iron sequestration (via GFP-based genetic iron sensor) and magnetic column retention relative to overexpressing the wildtype ftnA.

### Magnetic Column Retention characterization

A high-gradient magnetic column (Miltenyi LD columns) was sandwiched between two neodymium permanent magnets (K&J Magnetics Inc., BX8C4-N52) to create high magnetic field gradients inside the column. The column is first wetted by passage of 2ml of PBS 1X buffer. Then 500μl of cells re-suspended in PBS 1X buffer were added and flowed through by gravity into the elution tube, followed by addition of 3ml of PBS 1X buffer to wash through any unbound cells into the elution tube. Once dry, the column is removed from the permanent magnets, and 3ml of PBS 1X buffer is pushed through the column to extract the magnetically retained cells into a separate retention tube. Measuring OD600 of the elution and retention tubes allow estimation of cell counts and the percentage of total cells retained by the magnetic column.

### Iron level characterization by genetic sensor

For the genetic iron sensor, the *E. coli* fiu promoter was cloned along with a super-folder GFP (sfGFP) reporter via Gibson Assembly into a low copy (p15A origin), chloramphenicol-resistance plasmid compatible with the ferritin-expressing plasmid. Iron levels were measured for cells containing the ferritin-expression and iron sensor plasmids by taking the GFP fluorescence of the culture of cells (488 nm excitation by laser, 512 nm emission) in 96-well plate format using the BioTek NEO plate-reader. For calibration, known concentrations of iron sequesterer bi-pyridine were added to cell cultures. The fluorescence measured were normalized to culture density by dividing by OD600 measured by the same plate-reader. The increase in normalized fluorescence of the cells was plotted against the increase in bipyridine (or consequent decrease in free iron) and modeled (Fig S2) to determine the conversion between fluorescence reading and free iron concentration.

### Iron level characterization by Inductively-Coupled Plasma Mass Spectrometry (ICP-MS)

Cells to be measured were resuspended in 20% ultrapure nitric acid and mixed overnight (>12hrs) to lyse and digest organic material. The solution is then fed to Perkin Elmer 6100 ICP-MS machine (Trace Metal Lab, Harvard School of Public Health) to determine the concentration of a variety of elements (Fe, Co, Ni, Zn, Cd, As, Mg, Mn, U, Li) Prior to sample measurement, machine calibration was performed using solutions containing a linear range of known concentrations of the measured metals.

### Magnetic Resonance Imaging (MRI) characterization

MRI characterization was performed on a Brucker 9.4T MRI system at the MRI preclinical core of the Beth Israel Deaconess Medical Center. Cells were first washed of metal in culture and resuspended in metal-free PBS 1X buffer and subsequently loaded into NMR tubes. The NMR tubes were further immersed in DI water in a container to prevent artefacts due to susceptibility mismatch at the glass/air interface. Once samples were loaded into the MRI system, T2^*^ weighted relaxation times were obtained from fits of the signal decay after a gradient-echo pulse sequence and compared among samples.

### Superconducting Quantum Interference Device (SQUID) Magnetometry

Cells were first washed by DI water three times to remove excess iron, then pelleted by high-speed centrifuge (4000g). The cell pellets were flash-frozen by dipping into liquid nitrogen, followed by sublimation in a lyophilizer for one hour to remove water. The desiccated pellets were weighed and loaded into the Quantum Design MPMS SQUID and the sample magnetization (moment) was measured as magnetic field was varied in steps.

### Imaging *E. coli* culture over permanent magnets

*E. coli* cells induced to express ferritins were grown to saturation in LB media supplemented with 100μM iron and back diluted 1:5 with fresh LB media with 20% OptiPrep Density Gradient Medium to help suspend the cells. The cultures were dispensed into wells of a 6-well titer plate, where ring magnets were placed directly under the bottom of the wells, and left undisturbed. Plates were imaged by a digital camera after overnight growth to capture visible aggregation of cells toward the magnets.

### TEM of *E. coli* cross sections

*E. coli* cells in culture were resuspended in PBS 1X and fixed with glutaraldehyde overnight. The samples were subsequently dried, embedded in resin, and sectioned without additional heavy metal staining. The sections were imaged using the Tecnai G^2^ Spirit BioTWN microscope at variable acceleration voltage and magnification to maximize contrast of the electron dense nanoparticles in the cell cross-sections.

### SDS gel analysis of protein expression levels

*E. coli* cells were resuspended in SDS Buffer (NuPAGE LDS Buffer) with reducing agent, followed by two cycles of boiling at 95°C for 5 minutes and vigorous vortexing to lyse cells and denature proteins. The lysate was centrifuged to pellet cell debris, and the protein suspension was diluted and added to NuPAGE 4−12% Bis-Tris gel with MES buffer. For calibration of protein concentration via densitometry analysis, known dilutions of BSA protein in the same buffer and similarly denatured were added to the same gel containing the samples. Empty lanes in the gel were filled with equal volume of SDS buffer. After running at 200V for 35 minutes, the gel was removed and stained with Coomassie Orange dye for one hour and subsequently imaged for dye fluorescence on a Typhoon Imager. Densitometry analysis of the gel bands was conducted using ImageJ.

## Acknowledgements

We thank Professor Aaron Grant and Dr. Gopal Varma of the Beth Israel Deaconess Medical Center for assistance with MRI. We thank Maria Ericsson and the staff at Electron Microscopy Facility at Harvard Medical School for assistance with electron microscopy. We thank Thomas Ferrante and Garry Cuneo of Wyss Institute for experimental assistance. We thank Professor Ronald Walsworth, Dr. Pauli Kehayias, Dr. Cameron Myhrvold and Chenchen Luo for general discussions.

## Author contributions

X.L. designed the research, performed the experiments, analyzed the data and wrote the paper. P.L. performed magnetic evolution experiment and bioinformatics analysis. T.G. M.G., J.W., and P.S., analyzed the data and contributed to experiment design. T.G. and P.S. edited the paper.

## Competing financial interests

The authors declare no competing financial interests.

